# Requirements for efficient cotranscriptional regulatory switching in designed variants of the *Bacillus subtilis pbuE* adenine-responsive riboswitch

**DOI:** 10.1101/372573

**Authors:** Lea K. Drogalis, Robert T. Batey

**Affiliations:** Department of Biochemistry, University of Colorado, Boulder, CO 80303, USA

## Abstract

Riboswitches, generally located in the 5’-leader of bacterial mRNAs, direct expression via a small molecule-dependent structural switch informing the transcriptional or translational machinery. While the structure and function of riboswitch effector-binding (aptamer) domains have been intensely studied, only recently have the requirements for efficient linkage between small molecule binding and the structural switch in the cellular and cotranscriptional context begun to be actively explored. To address this, we have performed a structure-guided mutagenic analysis of the *B. subtilis pbuE* adenine-responsive riboswitch, one of the simplest riboswitches containing a secondary structural switch. Using a cell-based fluorescent protein reporter assay to assess ligand-dependent regulatory activity in *E. coli*, these studies revealed previously unrecognized features of the riboswitch. Most importantly, it was found that local and long-range conformational dynamics in two regions of the aptamer domain have a significant effect upon efficient regulatory switching. Further, sequence features of the expression platform including the pre-aptamer leader sequence, a nucleation helix and a putative programmed pause have clear affects upon ligand-dependent regulation. Together, these data point to sequence and structural features distributed throughout the riboswitch required to strike a balance between rates of ligand binding, transcription and secondary structural switching via a strand exchange mechanism.

## Introduction

Riboswitches are RNA elements that can adopt two distinct structures to direct expression of a message, depending upon the occupancy status of a small-molecule receptor domain (known as the aptamer domain) [1-3]. These structures generally inform transcription through differential formation of a rho-independent transcriptional terminator or translation by either occluding or exposing the ribosome binding site, although a diverse set of other mechanisms of expression regulation have been found [4-6]. This straightforward mechanism of ligand-dependent control of gene expression without the assistance of accessory proteins has made these devices attractive for use in various synthetic biology applications [7, 8]. In part, the means to raise RNA sequences that bind with high affinity and specificity to a broad spectrum of small molecules using *in vitro* selection approaches and the modularity of RNA secondary structure make the design and implementation of such input-output devices conceptually simple [9, 10].

However, engineering modular riboswitches and other RNA devices has been problematic, such that only a few robust regulatory elements have achieved real-world application [11]. This suggests that the basic understanding of how these RNAs mechanistically translate ligand occupancy of the aptamer domain into an observable response remains incomplete. One major hurdle in the robust design of riboswitches is that they generally only function in the context of transcription [1, 12-14], requiring careful consideration of kinetic processes such as rates of ligand binding, higher-order RNA folding and secondary structural switching processes such as strand exchange [15]. Furthermore, the influence of RNA sequence on processes such as transcriptional pausing and participation of factors such as NusA or Rho on riboswitches in the cellular context further confounds their design and implementation [16-18]. It is likely that many of these processes are influenced by cryptic sequence elements difficult to engineer into an RNA of interest.

Natural riboswitches represent important model systems for understanding the sequence features that enable how RNA devices can be engineered to be efficient ligand-dependent genetic regulators that function within a broad spectrum of industrially useful bacteria [19]. One of the simplest riboswitches containing a discrete ligand-binding domain and secondary structural switching domain (otherwise known as the expression platform) is the *Bacillus subtilis* adenine-responsive *pbuE* regulatory element (Fig 1) [20], making it ideal for detailed analysis of coupling of ligand binding to regulatory activity [21]. The aptamer domain shares the same general three-dimensional architecture as all purine riboswitches with the ligand binding site embedded within the three-way junction, adjacent to the 3’-side of the P1 helix [22, 23]. Overall organization of the aptamer is achieved by a loop-loop interaction (L2 and L3, Fig 1) distal to the junction and P1 helix whose formation is ligand-independent in some riboswitches such as the *B. subtilis xpt-pbuX* guanine-sensing riboswitch [24-26] and dependent on ligand binding in others such as the *pbuE* aptamer [27-29]. When ligand is bound, the aptamer including the P1 helix is stabilized against formation of an alternative structure enabling full transcription of the mRNA and expression of the encoded purine efflux pump under high adenine conditions [20].

**Fig 1.**
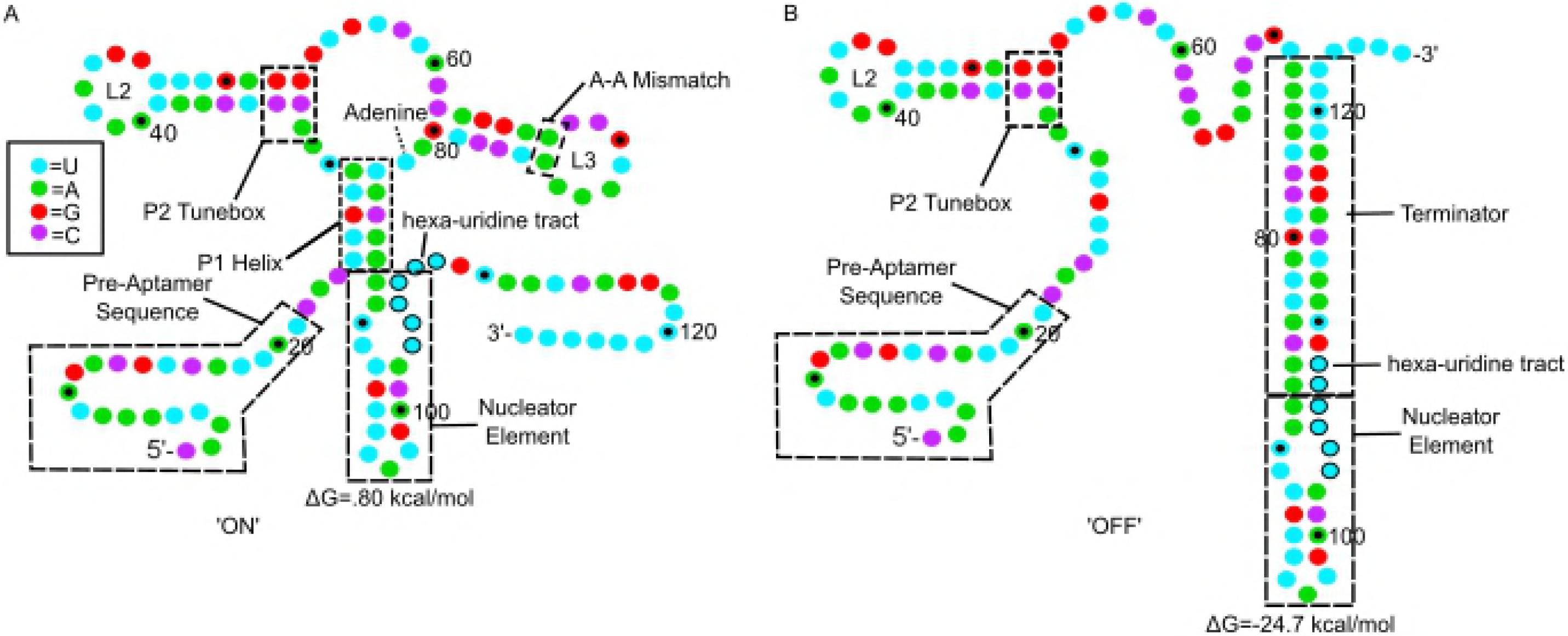
*pbuE* secondary structures. (A) Secondary structure of the *B. subtilis* adenine-responsive *pbuE* riboswitch starting at the transcriptional start site and ending at the poly-uridine tract in the ligand-bound state (ON). Relevant structural elements include the pre-aptamer leader sequence, P1 helix, P2 tunebox, P3 A-A mismatch and the P2-P3 loop-loop interaction. In this scheme, cyan represents uridine, green represents adenine, red represents guanine, and purple represents cytosine; residue numbering reflects the experimentally determined start site of transcription [30]. (B) Secondary structure of the *pbuE* riboswitch in the unbound state (OFF) with the intrinsic terminator formed at the expense of the P1 and P3 helices.

Single molecule analysis of the adenine-dependent folding of the *pbuE* riboswitch in the context of transcription has suggested a mechanism for how ligand binding to the aptamer domain prevents formation of the competing intrinsic terminator element [31]. In this model, during transcription the aptamer hierarchically and rapidly folds into a productive conformation by the time the polymerase reaches a hexa-uridine tract found at the 3’-side of the nucleator element. The nucleator element caps the terminator element and its structure is independent of ligand binding (Fig 1). Shortly thereafter, sequence on the 3’-side of the helix begins to strand exchange through the P1 helix. In the absence of ligand, this migration rapidly continues through J3/1 and P3 to form the functional terminator helix that prompts the RNA polymerase to disengage from RNA synthesis, and thereby gene expression is repressed. However, in the presence of ligand, strand exchange is impeded by ligand-dependent organization of the three-way junction and formation of base triples involving J2/3 and the two junction-proximal base pairs of P1. This set of triples is proposed to act as a kinetic roadblock to further strand exchange into the junction and P3, providing the RNA polymerase sufficient time to synthesize the message past the intrinsic terminator’s poly-uridine tract and escape the riboswitch. Together with other biochemical and biophysical studies, there is compelling evidence that the *pbuE* riboswitch is a “kinetically controlled” switch such that it does not reach thermodynamic equilibrium with respect to adenine binding during the timeframe of transcription of the leader sequence [12, 31]. Ultimately, the terminator form of the riboswitch represents the most thermodynamically stable state of RNA and the riboswitch adopts this structure regardless of ligand concentration [32].

Despite serving as a model system for understanding the physical basis for the coupling of binding and regulation, the *pbuE* riboswitch has not been extensively investigated in the cellular context. Because many riboswitches only function in the context of transcription, understanding the sequence and structural requirements of the expression platform conferring ligand-dependent activity requires a cell-based reporter assay. Towards this end, a robust fluorescent protein reporter assay has been established for the *pbuE* riboswitch using the adenine analog 2-aminopurine (2AP) as the ligand [33]. This study revealed several facets of the expression platform that influence ligand-dependent regulatory activity. The most important finding was that the riboswitch is highly tolerant to significant variation in the length of the P1 helix. This contrasts results of biophysical studies which proposed that P1 helix length and associated thermodynamic stability are critically important for proper switching. Instead, this result supports a low-energy strand displacement mechanism that governs formation of the riboswitch’s alternative secondary structures [33]. Second, these data suggested that a ligand-independent secondary structure in the expression platform that we call the “nucleator element” plays an important role in regulatory activity. This indicates that sequences and secondary structures within the expression platform influence the regulatory response.

In the current study, we seek to further define regions of the *pbuE* riboswitch that influence the magnitude of the ligand-dependent regulatory activity using a structure-guided mutagenic approach. To assess whether the aptamer domain has features beyond the conserved ligand binding site that are required for efficient secondary structural switching, a set of *B. subtilis* purine-responsive aptamers was spliced into the *pbuE* expression platform to create chimeras homologous to the native *pbuE* riboswitch. Despite very high sequence and structural similarities, these designed chimeras fail to function in *E. coli*. Regulatory activity is restored by introducing point mutations in two regions that influence the conformational ensemble of the unliganded state, suggesting new roles for sequences in the aptamer domain. To further clarify structural effects on strand invasion dynamics, we examined the sequence of the P1 helix, the nucleator element, and a putative programmed pause. These mutants revealed altered regulatory properties that indicate the nucleator element and an adjacent six uridine tract are crucial for efficient ligand-dependent upregulation, while the sequence of the P1 helix is likely tuned to influence the rate of strand invasion immediately prior to the key regulatory roadblock. Together, these data reveal new aspects of how the riboswitch’s activity is modulated by key sequence and structural elements and suggest new design strategies for the engineering of novel riboswitches.

## Materials and methods

### Construction of reporter plasmids

For each riboswitch a set of overlapping DNA oligonucleotides was synthesized to construct the gene using recursive PCR [34]. Using standard molecular cloning techniques [35], each riboswitch variant (S1 Table) was cloned upstream of the *gfpuv* gene to regulate its expression in a ligand-dependent manner. The parental vector containing *gfpuv* is derived from a low-copy pBR327 plasmid [36] and contains the strong *rrnB* terminator upstream of a synthetic insulated promoter [37] of moderate strength to limit transcription of upstream genes. S1 Table provides full verified sequences of each riboswitch from the first transcribed nucleotide to the poly-uridine tract.

### Cell-based fluorescence assays

*E. coli* K12 strain BW25113 (Keio knockout collection parental strain) [38] was transformed with sequence-verified reporter plasmid using standard protocols [35]. Transformants were plated on 2xYT growth medium agar plates containing 100 μg/mL carbenicillin for resistance marker selection. A single colony was picked and used to inoculate an overnight 2 mL culture of 2xYT containing 100 μg/mL of ampicillin and grown at 37 °C. A 20 μL volume of overnight culture was used to inoculate three individual 2 mL cultures of rich, chemically defined growth (CSB) medium [36] containing 100 μg/mL of ampicillin. Three additional 2 mL CSB plus ampicillin cultures containing 1 mM 2-aminopurine were also inoculated with saturated overnight cultures in a 1:100 dilution. These secondary cultures were grown to an OD_600_ between 0.2-0.4 at 37 °C. Expression was determined by collecting the OD_600_ (cell density) and fluorescence intensity of 175 μL of each culture using an Infinite M200 PRO plate reader (Tecan). Cells were excited at 395 nm and read at 510 nm for all fluorescence measurements with maximum fluorescence being set with a well containing 175 μL of 3 μM fluorescein standard. Normalized fluorescence per cell density was calculated by dividing fluorescence measurements by the corresponding cell density values. Fold induction was calculated by subtracting background fluorescence (normalized fluorescence of cells containing only parental vector) from normalized fluorescence values and fluorescence values of cultures containing ligand divided by those without ligand. Data for each biological replicate were collected on different days to account for day-to-day variation; all data is based upon three technical repeats of three independent biological replicates (S2 Table).

### Co-transcriptional RNA folding simulations

The co-transcriptional folding landscapes of select mutant riboswitches were modeled using Kinefold (http://kinefold.curie.fr; [39]) Riboswitch RNA sequences starting at the transcription initiation site and ending after the poly-uridine tract of the rho-independent transcriptional terminator were used in the simulation with the transcriptional speed set as a new nucleotide (nt) added every 20 milliseconds (50 nt sec-1) with pseudoknots disallowed. Helical tracing graphs were obtained from these simulations as well as folding movies showing the lowest free energy structure with the addition of each base. The resultant folding trajectories allow for visual determination of persisting secondary structures that are not reported in the helical folding graphs.

## Results

### Other purine riboswitch aptamer domains can drive the *pbuE* expression platform

It has been previously hypothesized that expression platforms are modular and capable of hosting different aptamers to create “chimeric” riboswitches, which has been shown experimentally for both “OFF” and “ON” switches using a single-turnover *in vitro* transcription assay [11, 36, 40]. To determine whether the *pbuE* structural switch can be regulated by other purine riboswitch aptamers, a conservative chimeric riboswitch was designed that fused the *B. subtilis xpt-pbuX* guanine/hypoxanthine-responsive riboswitch aptamer domain [22] with the *B. subtilis pbuE* adenine-responsive riboswitch expression platform (Fig 2a). For these studies, the C74U variant of the *xpt* aptamer was used that switches its specificity to adenine/2-aminopurine, which has been extensively used to study this aptamer and does not alter the properties of this RNA [41-43]. Note that while only P2 and P3 are boxed in Fig 2a, the junction-proximal two base pairs of P1 as well as all nucleotides in J1/2 and J3/1 are identical between the *xpt*(C74U) and *pbuE* aptamers as well as seven out of eight nucleotides in J2/3. The only nucleotide difference in J2/3 is G56 of *pbuE*, the equivalent position being uridine in *xpt*. Thus, the primary differences in these aptamer domains lie in P2/L2 and P3/L3. Since the P3 sequence is different between these two aptamers, compensatory changes were made in the expression platform of the *xpt*(C74U)/*pbuE* chimera to ensure formation of a stable transcriptional terminator hairpin that is the same length as that of the wild type *pbuE* riboswitch (Fig 2a). As the secondary and tertiary structure of all purine riboswitches is highly conserved, we hypothesized that this conservative aptamer substitution in the chimera (*xpt/pbuE* A) would robustly turn on reporter protein expression in the presence of 2-aminopurine (2AP). However, compared to the wild type *pbuE* riboswitch, the chimera exhibited no activation of gene expression, with low reporter expression in both the presence and absence of 2AP indicating strong constitutive termination (Fig 2B).

**Figure 2.**
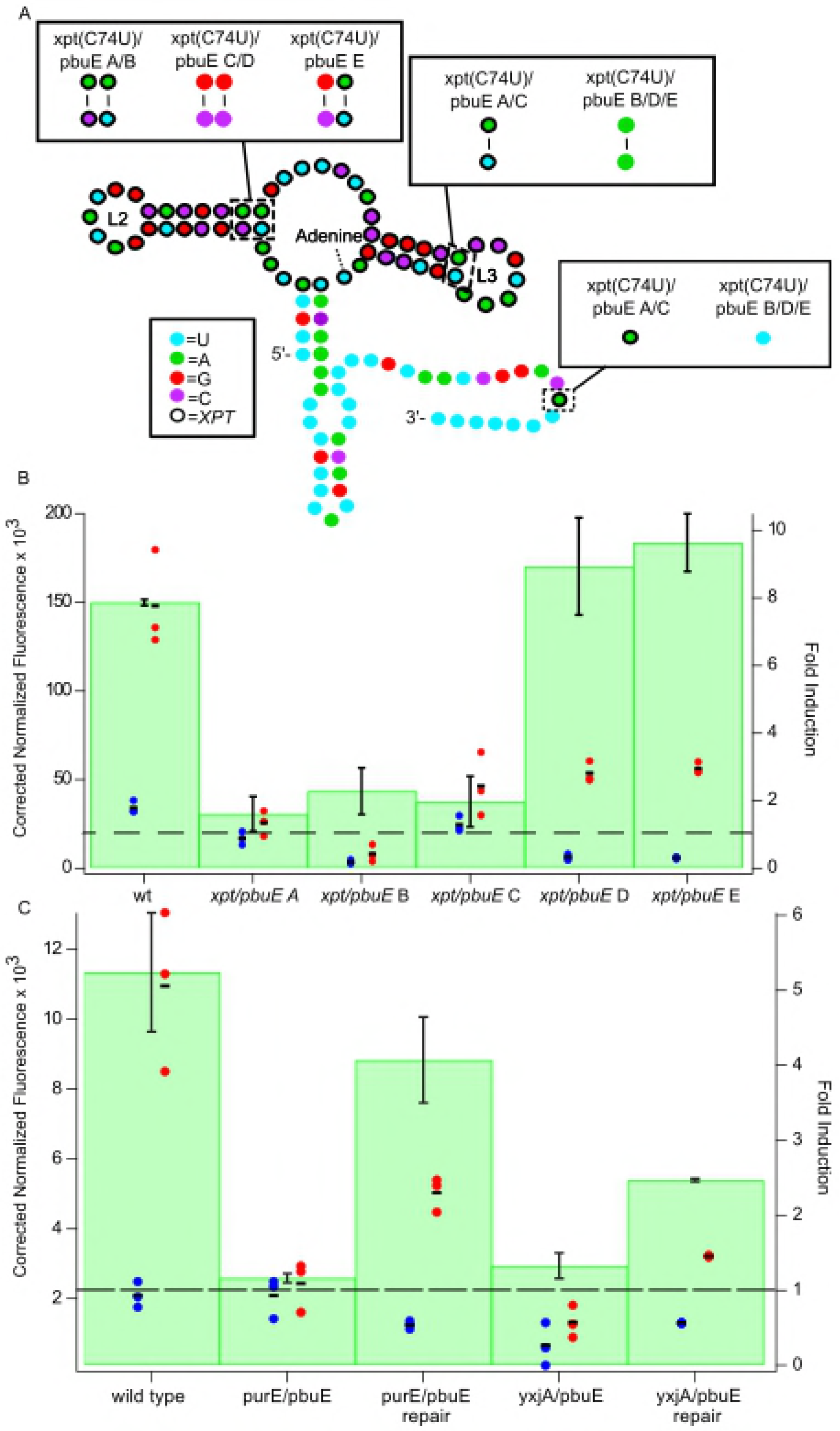
Multiple mutations are required for successful switching of the *xpt*(C74U)/*pbuE* chimera. (A) Secondary structure of the *xpt*(C74U) aptamer appended to the expression platform of the *pbuE* riboswitch. The pre-aptamer sequence has been omitted for clarity. Relevant structural elements include the P2 tunebox and the P3 distal A-A mismatch. The nucleotide coloring scheme is the same as in Fig 1; residues circled in black are part of the *xpt*(C74U) aptamer. (B) Normalized fluorescence data for each *xpt*(C74U)/*pbuE* chimera variant in the absence (blue) and presence (red) of 2-aminopurine. Each data point represents the average of three technical replicates of one biological replicate; the black bar represents the average of the biological replicates. Green bars represent fold induction. (C) Normalized fluorescence data for *purE*(C74U)/*pbuE* and *yxjA*(C74U)/*pbuE* chimera variants in the absence (blue) and presence (red) of 2-aminopurine.

To understand why this chimeric RNA fails to regulate gene expression, an extensive mutational analysis of the aptamer domain was performed in which elements of the *xpt*(C74U) P2 and P3 were systematically substituted back for the *pbuE* sequences. While most of these variants failed to rescue regulatory activity (data not shown), it was found that two specific point substitutions made in the *xpt*(C74U) aptamer resulted in robust 2AP-dependent gene regulation (Fig 2B). The first point substitution was in a region of the aptamer referred to as the P2 “tune box”, which comprises the junction-proximal two base pairs in P2 helix along with the last nucleotide of J1/2 (Fig 1A) [43]. Natural variation in this region of the purine riboswitch aptamer has a significant effect on its kinetic and thermodynamic ligand binding properties. It was hypothesized based upon chemical probing data that this region affects the organization of the three-way junction in the absence of effector ligand [43]. However, introducing a tune box sequence identical to that of *pbuE* into the *xpt*(C74U)*/pbuE* chimera (*xpt/pbuE* C) alone was insufficient to restore switching ability (Fig 2B).

The second region that affects regulatory activity of the *xpt*(C74U)*/pbuE* chimera is the base pair in P3 proximal to L3. To achieve the “OFF” state, the *pbuE* terminator must invade into L3 and disrupt the stable L2-L3 interaction in the process. Single molecule and fluorescence lifetime studies demonstrated that while the *xpt* L2-L3 interaction is stable in the absence of ligand [25, 26, 29]; the equivalent interaction in *pbuE* is only transiently stable in the absence of ligand binding [27, 28]. Further, introduction of an A-A mismatch as seen in *pbuE* at the base of the *xpt* L3 destablizes the L2-L3 interaction [24]. Introducing a point substitution into the *xpt*(C74U)/*pbuE* hybrid to yield an A-A mismatch (*xpt/pbuE* B) was insufficient to restore regulation (Fig 2B). In contrast, simultaneously introducing the *pbuE* P2 tune box and L2-L3 destabilization element into the hybrid (*xpt/pbuE* D) successfully restored switching activity to that of wild type with decreased leaky gene expression in the absence of ligand as compared to wild type (Fig 2B). Furthermore, simply repairing the mismatched base pair in the *xpt* aptamer domain P2 to a Watson-Crick (*xpt/pbuE* E) also restores switching performance and improves leakiness when introduced in concert with the distal A-A mismatch. These data strongly suggest the conformational dynamics of the aptamer and associated crosstalk between the L2-L3 interaction and the three-way junction in the unliganded state are critical for establishing a strong ligand-dependent regulatory response.

To determine whether these two features of the *xpt* aptamer also influence the ability of other guanine-binding aptamers to regulate the *pbuE* structural switch, we investigated several other chimeras. Using the same strategy as with *xpt* to create a 2AP-responsive chimera, the *B. subtilis yxjA* (S5 Fig) and *purE* (S4 Fig) guanine riboswitch aptamer domains [44] were placed into the wild type *pbuE* context. The wild type *purE* riboswitch aptamer domain lacks both the P3 A-A mismatch and a fully base paired P2 tune box, leading to the design of a “repaired” *purE* with the same mutations as the optimal *xpt/pbuE* E construct (*xpt* A-A P3 plus paired tune box). Consistent with *xpt*(C74U)/*pbuE*, the *purE*(C74U)/*pbuE* hybrid did not show any switching activity (Fig 2C) and gene expression in the presence or absence of 2AP was low. The repaired *purE*(C74U)/*pbuE* hybrid showed improved termination in the absence of 2AP and increased aptamer stability in the presence of 2AP with a 4-fold induction of reporter expression (Fig 2C). Switching performance was not as effectively repaired in the case of *purE*(C74U)-*pbuE* compared to *xpt*(C74U)*-pbuE,* but a significant restoration of regulatory activity was still observed.

Unlike the *purE* guanine-sensing aptamer, the wild type *yxjA* aptamer contains a P3 A-A mismatch at the base of L3 as well as a P2 tune box with two Watson-Crick pairs like the *pbuE* aptamer. Thus, this aptamer was anticipated to work with the *pbuE* expression platform and yield effective switching without altering its sequence. Surprisingly, the yxjA(C74U)/*pbuE* chimera displayed little 2AP-dependent induction of expression (Fig 2C). A variant of *yxjA* containing the tune box sequence identical to the *pbuE* aptamer was also examined; this repaired *yxjA*(C74U)/*pbuE* chimera showed only modest switching activity, though expression increased in both the presence and absence of 2AP relative to the unrepaired chimera. Together, these data suggest that two elements within the P2 and P3 helices of the purine aptamer domain that often harbor base mismatches, while not directly involved in ligand binding, are nonetheless critical for establishing a regulatory response. However, it is clear there remain other still unidentified features in the aptamer domain that are important for regulatory activity.

### The P1 helix has a modest influence on regulatory activity

The above data reveal that sequence features in P2 and P3 have a direct impact upon regulatory activity. It is also likely that the sequence of the P1 helix, a critical player in the regulatory switch, has a significant role in promoting the ligand-dependent structural switch. In a prior study, it was hypothesized that the sole G-C base pair in *pbuE* P1 represents an important road block to rapid invasion through P1, giving the aptamer sufficient time to bind effector [45]. To directly test this hypothesis, mutations were made to P1 that either replace the G-C pair with an A-U pair or add G-C pairs to the junction-distal region of the P1 helix (Fig 3A). Mutations designed to weaken the P1 helix were anticipated to decrease expression in the absence of ligand due to the increased ease of strand invasion by the expression platform through P1 while expression in the presence of ligand would decrease due to destabilized aptamer folding. When the P1 helix is weakened with an A-U Watson-Crick or G-U wobble pair (P1-AU, P1-GU), a small increase in fluorescence relative to wild type was observed in the absence of ligand along with more significant decreases in fluorescence relative to wild-type in the presence of ligand, in support of this hypothesis (Fig 3B).

Conversely, adding G-C pairs to P1 was expected to decrease expression in the absence of ligand as strand invasion through P1 is impaired. Given this expectation, it was unanticipated that increasing the G-C base pair content of the P1 helix with G-C pairs (P1-GC2a, -GC2b and -GC3) results in only slightly less expression in the absence of ligand than wild-type while expression in the presence of ligand gradually decreased. The most marked decrease in regulatory activity occurs in the P1-GC3 variant, whose levels of expression are low regardless of 2-AP and overall fold-induction is below 2.

**Fig 3.**
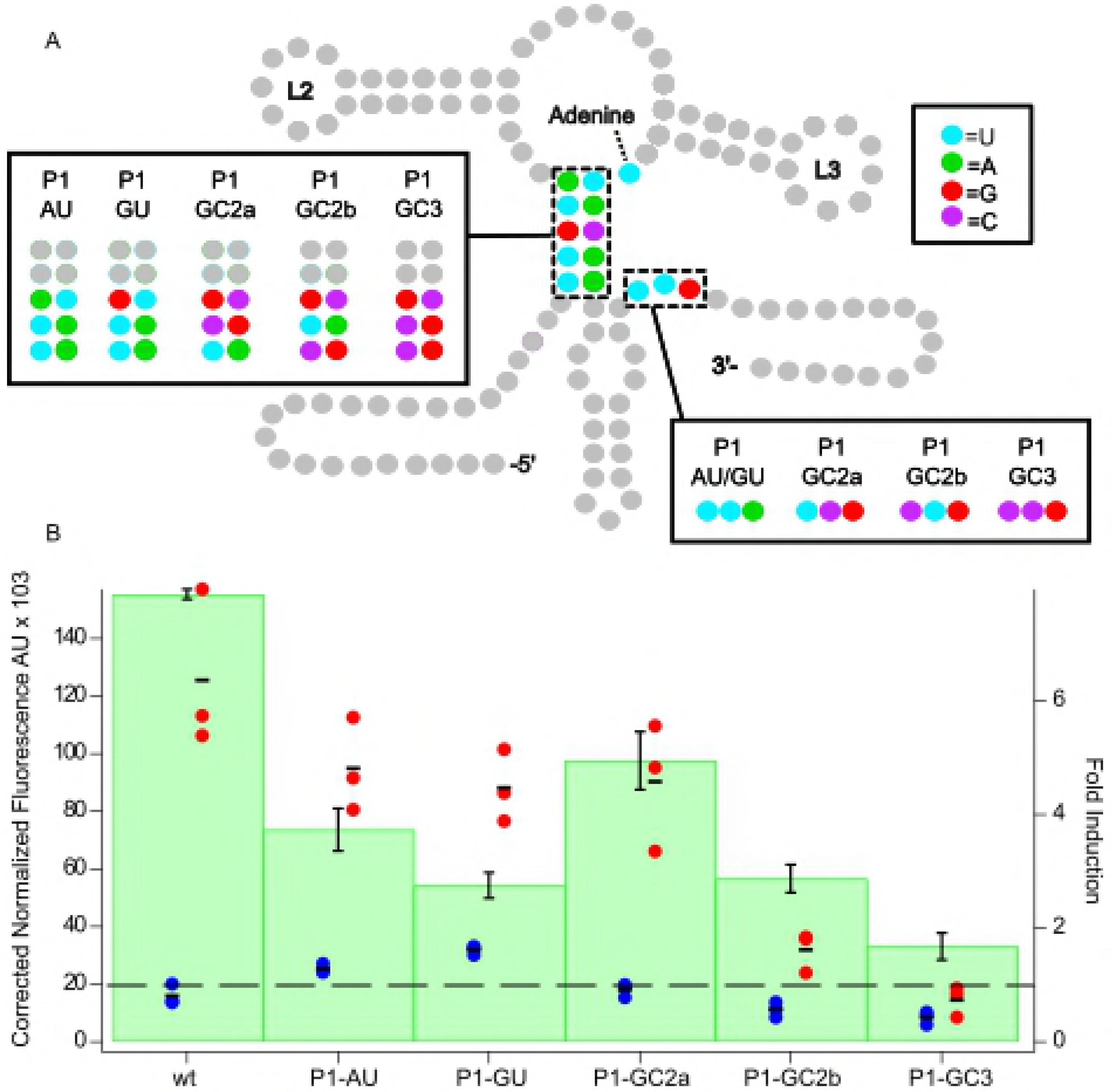
Altering P1 sequence impacts regulatory activity. (A) Secondary structure of the *pbuE* riboswitch highlighting each mutant of the P1 helix. Relevant structural elements include the P1 helix and the P1 helix pairing region of the expression platform. The nucleotide coloring scheme is the same as in Fig 1. (B) Normalized fluorescence data for each P1 helix mutant in the absence (blue) and presence (red) of 2-aminopurine. Each data point represents the average of three technical replicates of one biological replicate. The black bar represents the average of the biological replicates; green bars represent fold induction. pBR327 is a negative control.

### Altering the pre-aptamer leader sequence unmasks a strong influence of P1 sequence on regulatory activity

While the above results reveal a modest influence of the sequence composition of the P1 helix on activity, the effects of overall trends were not clear. In particular, it was surprising that P1-GC2b and -GC3 did not show strong expression of the reporter in the absence of ligand, suggesting that the aptamer domain was somehow debilitated despite stabilizing the P1 helix. One possibility is that mutations in P1 resulted in alternative structure(s) that disrupt formation of a functional aptamer or terminator. To explore this, a subset of sequences were analyzed by Kinefold, a program that simulates co-transcriptional folding pathways [39]. Analysis of the wild type *pbuE* riboswitch used in this study reveals a folding pathway in which P2 and P3 fold, followed by a brief formation of P1 that is then disrupted by folding of the terminator--consistent with experimentally derived models of the *pbuE* folding pathway ([45], [31]). However, in both the P1-GC2b and P1-GC3 variants, the helical formation traces revealed the appearance of alternate pairing schemes involving the 5’-side of P1 and the pre-aptamer leader sequence, indicating alternative folds disruptive to the P1 helix and aptamer formation.

With this hypothesis in mind, these two variants were redesigned to minimize alternative pairing. Using Kinefold simulations, the pre-aptamer sequence was altered to prevent alternative pairing with P1 helical elements (P1-GCb,rep and P1-GC3,rep). These “repair” variants (Fig 4A) show different regulatory properties. Repairing the mismatched folding in P1-GC2b was sufficient to restore switching to the level of wild type *pbuE*. Notably, only a few point substitutions of the pre-aptamer sequence that destabilize an alternative helix are sufficient to restore activity (Fig 4B). Conversely, repair of P1-GC3 either by point substitutions or a larger deletion did not greatly improve switching.

**Figure 4.**
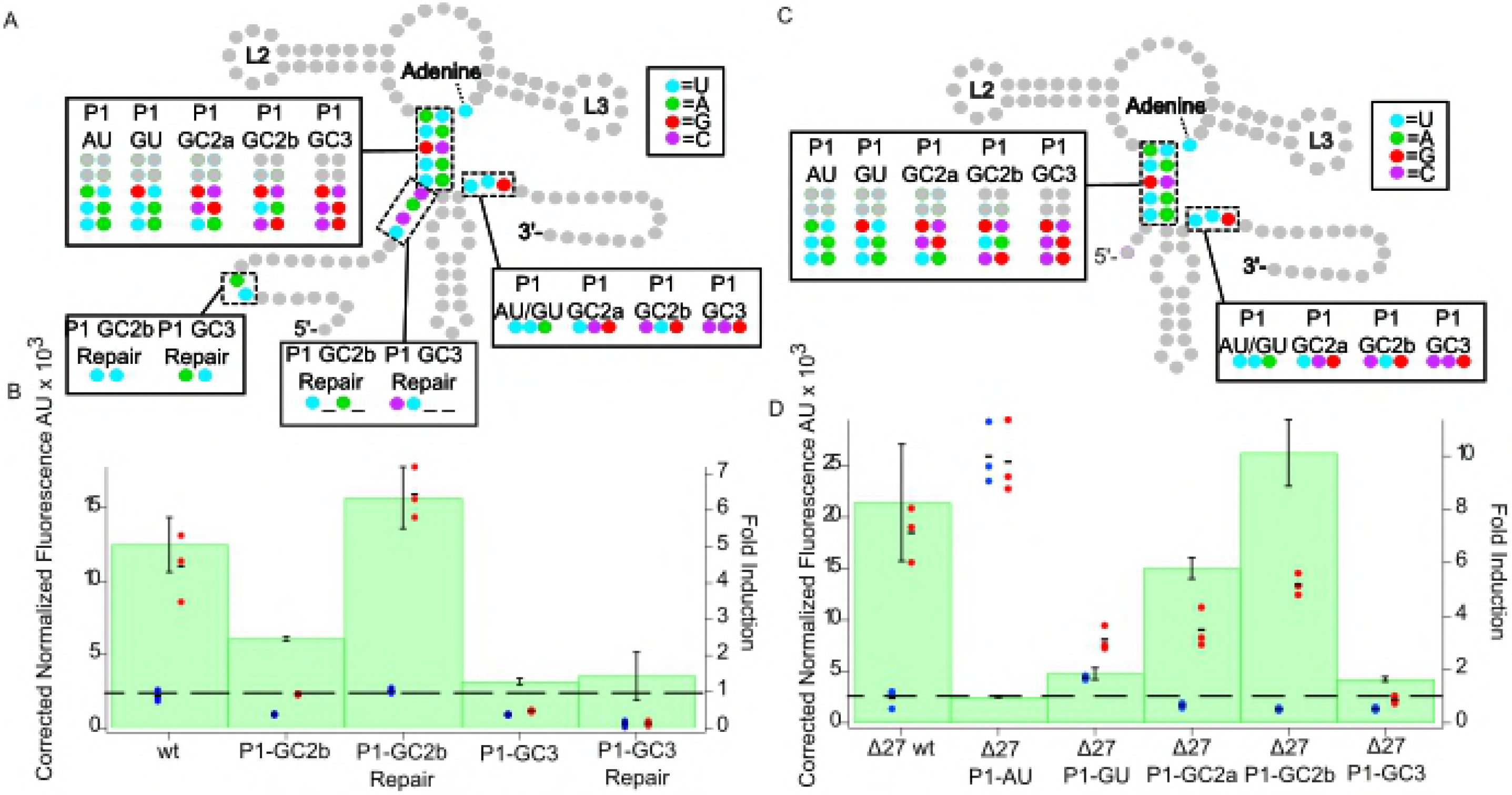
Repair and removal of the pre-aptamer sequence helps clarify trends of P1 variants. (A) Secondary structure of the *pbuE* riboswitch including mutations in the leader sequence to ablate unintended pairing interactions. Mutations in the P1 helix are highlighted as are the compensatory changes required in the expression platform. The nucleotide coloring scheme is the same as in Figure 1. (B) Normalized fluorescence data for P1-GC2b and P1-GC3 mutants without the leader sequence or with a repair in the absence (blue) and presence (red) of 2-aminopurine. Each data point represents the average of three technical replicates of one biological replicate. The black bar represents the average of the biological replicates. Bars represent fold induction. (C) Secondary structure of the *pbuE* riboswitch without (∆27) the pre-aptamer sequence. Mutations in the P1 helix are highlighted as are the compensatory changes required in the expression platform. (D) Normalized fluorescence data for each P1 helix mutant without the leader sequence in the absence (blue) and presence (red) of 2-aminopurine.

The above data indicates that the pre-aptamer leader sequence can confound mutagenic analysis of riboswitch function via the formation of unanticipated alternative structure. To minimize this issue, a new set of P1 mutants was designed in which the pre-aptamer sequence was almost completely removed through a deletion of nucleotides 1-27 (Fig 1). This deletion in the context of the wild type riboswitch did not substantially alter its regulatory properties, although a small increase in the 2AP-dependent reporter gene expression was noted (S3 Fig).

In the absence of the pre-aptamer sequence (Fig 4C), a clear trend in the function of the P1 helix variations was revealed (Fig 4D). Variations that weakened the P1 helix, (∆27)P1-AU and (∆27)P1-GU showed no or very little ability to activate gene expression in the presence of 2AP. Notably, (∆27)P1-AU displayed a dramatic decrease in termination ability, suggesting that the terminator does not efficiently invade through the aptamer even in the absence of ligand binding. In contrast, (∆27)P1-GU is able to terminate, but not as well as other variants, suggesting that the weakened P1 helix (and weakened terminator helix in the complementary region) does not promote full strand exchange on the timescale that enables the regulatory switch. Variants that stabilized the P1 helix by adding another G-C base pair ((∆27)P1-GC2a and (∆27)P1-GC2b) display expression levels and fold-induction comparable to wild type, indicating that the riboswitch accommodates a more G-C rich P1 helix. However, further addition of a third G-C pair to the P1 helix ((∆27)P1-GC3) yields an RNA that is constitutively terminated, indicating that the G-C rich P1 helix likely supports highly efficient strand exchange to form the terminator helix.

### The nucleator element is critical for secondary structural switching

In a strand exchange model of ligand-dependent secondary structural switching, the terminator helix must rapidly nucleate to begin the process of strand exchange with the P1 helix of the aptamer domain. It has been observed that riboswitch expression platforms often contain a hairpin element that can be formed in either conformational state that is proposed to serve as a nucleation element (Fig 1A) for the secondary structure that competes with the P1 helix [33, 46]. Like related toehold-mediated strand displacement processes [47, 48], the kinetics of invasion is likely related to the length and/or base pair composition of the nucleator helix. To examine the importance of nucleator stem-loop length on regulatory activity, the wild type nucleator element was replaced with a set of designed nucleator helices of varying lengths capped with a stable GAAA tetraloop (Fig 5A). This series spans from 0 to 8 Waston-Crick base pairs in length (note that the wild type nucleator helix is 8 base pairs including two U-U mismatches), with their predicted free energies ranging from 0 to −17.2 kcal/mol.

**Figure 5.**
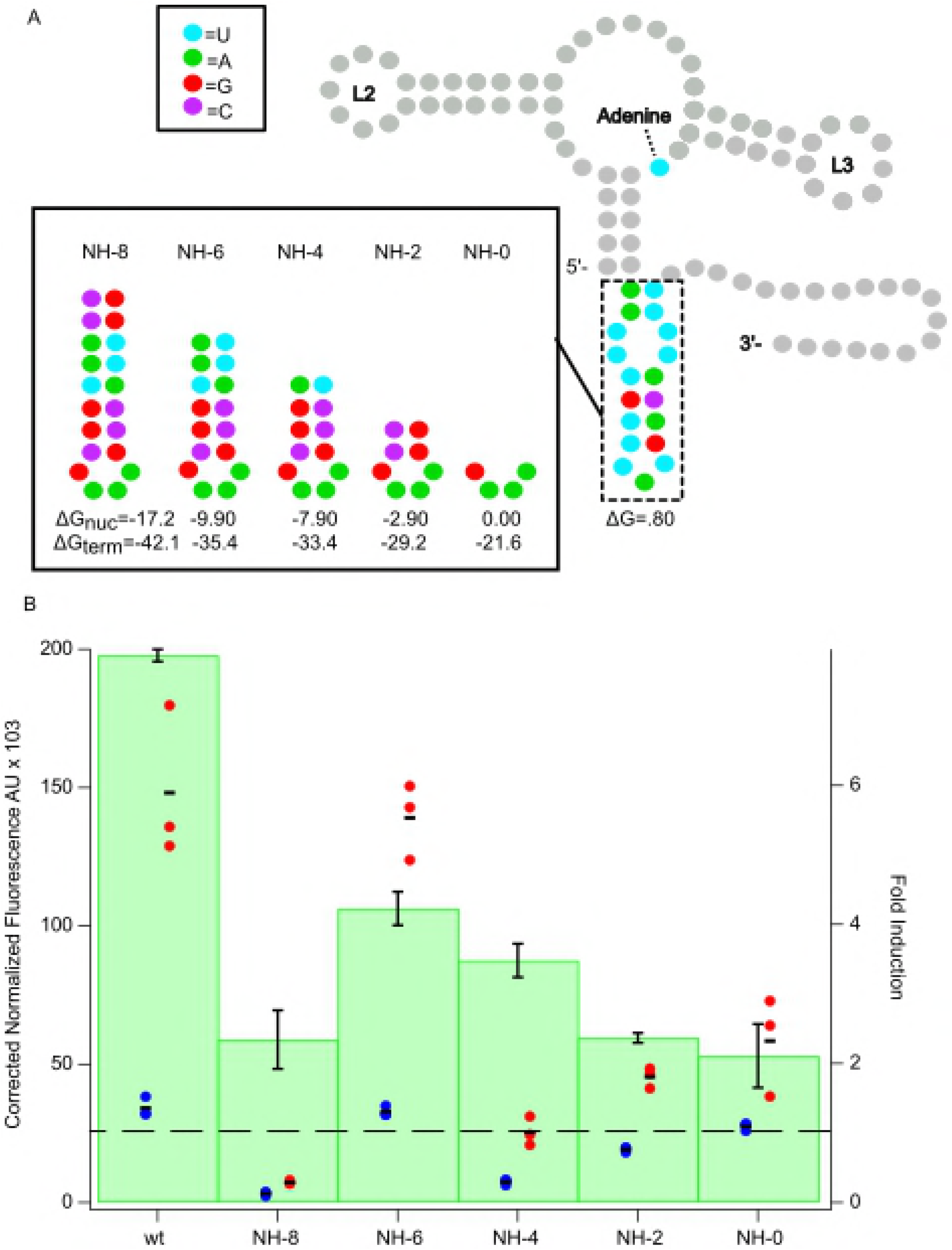
Nucleator element length impacts the efficiency of regulatory activity. (A) Secondary structure of the *pbuE* riboswitch highlighting each mutant of the nucleator element. The aptamer domain remains unchanged. The leader sequence has been omitted for simplicity. The nucleotide coloring scheme is the same as in Figure 1. (B) Background corrected normalized fluorescence data for each nucleator element mutant in the absence (blue) and presence (red) of 2-aminopurine. Each data point represents the average of three technical replicates of one biological replicate and the black bar represents fold induction of the average of the biological replicates. pBR327 is a negative control. Wild type data used for this figure collected on the same days as the other data in the figure.

This series of RNAs displays a trend towards increased fold induction in the presence of 2AP with one clear exception. (Fig 5B). As the length of the nucleator helix increases from 0 to 6 base pairs, the fold induction increases from 2- to over 4-fold, indicating that increasing helix length has a positive effect on the regulatory activity. NH-6 has the most similar expression levels of GFP reporter in the absence and presence of 2-AP to the wild type riboswitch, although the overall fold induction is lower. Further decreases in the length of the nucleator element yield systematically lower levels of fold induction (∼2-3-fold). Surprisingly, regulatory activity is not completely abolished in the NH-0 variant, indicating that the nucleator element is not essential for activating gene expression, albeit at low levels. Notably, the most conservative variant, NH-8, that preserves the overall length of the nucleator helix shows fold induction close to that of the 0 base pair variant, NH-0. This suggests that while the nucleator element’s length contributes to promoting ligand-dependent regulatory activity, its effect is potentially obscured by perturbing other key elements in this region of the expression platform.

A second feature of the nucleator element that may be important for efficient regulation is a potential “programmed pause” at the 3’-side of the helix. It well established that many riboswitches contain stretches of uridines in their expression platforms that represent RNA polymerase pause sites that give more time for events such as RNA folding and ligand binding to occur [13, 17, 49, 50]. In the *pbuE* riboswitch, a hexa-uridine tract is located at the 3’-side of the nucleator element that could serve as a pause site [13]. This uridine tract has been observed to be a pause site *in vitro* using *E. coli* and *B. subtilis* RNAP [30], but was not identified as a pause site in a genome-wide survey of transcriptional pausing in *B. subtilis* [51].

Other than wild type, none of the variants in the nucleation length series likely support transcriptional pausing at the uridine tract on the 3’-side of the nucleator element because the hexa-uridine tract at nucleotides 103-108 at the 3’-side of the nucleator helix have been altered. To determine the importance of this pause, a variant of NH-8 was tested in which two G-C pairs were converted to A-U pairs, thereby restoring the third and fourth uridines of the tract (NH-8 U) (Fig 6A). This change resulted in complete recovery of 2AP-dependent regulatory activity with a fold induction that exceeded wild type (Fig 6B). Without the pause site, it is likely that the terminator rapidly invades through P1 regardless of the presence of effector because the aptamer is not given sufficient time to bind ligand. If the stability of the nucleator helix were the only determinant of its influence on regulatory activity, then NH-8 should perform equally well as this variant. To confirm this hypothesis, a number of variants were designed that preserve the six uridines on the 3’-side, including a variant with a five base pair helix (NH-5 U) and variants of NH-4 and NH-0 from the original nucleator length series (NH-4 U, NH-0 U) (Fig 6C). NH-5 U was found to have better termination efficiency and switching ability than NH-6, NH-4, NH-2, NH-0 (Fig 6D) and NH-5 without the six uridine pause (Fig 6E), while NH-4 U and NH-0 U both outperform their hexa-uridine pause deficient counterparts, further suggesting that the uridine tract is critical for modulating the temporal window of transcription to enable key events more time to occur.

To explore whether the position of the hexa-uridine tract is crucial for efficient regulation, variants of NH-8, NH-8 U, and NH-5 containing a six uridine loop in place of the GNRA tetraloop were designed. In the case of NH-5 6U Loop, the performance of the riboswitch was equivalent to that of wild type and NH-5 (Fig 5G). NH-8 U 6U Loop (containing a hexauridine loop as well as a hexauridine tract) performed similarly to wild type but did not achieve the same performance as NH-8 U suggesting some impairment by the loop substitution. However, NH-8 6U Loop with only a hexauridine tract in the terminal loop (equivalent to NH-5 6U) displayed minimal switching ability (Fig 5D). Together, these data further indicate that the pause is important for aptamer binding rather than nucleation of the competing terminator element, however the exact mechanism of kinetic control via the hexauridine pause remains unclear.

**Fig 6.**
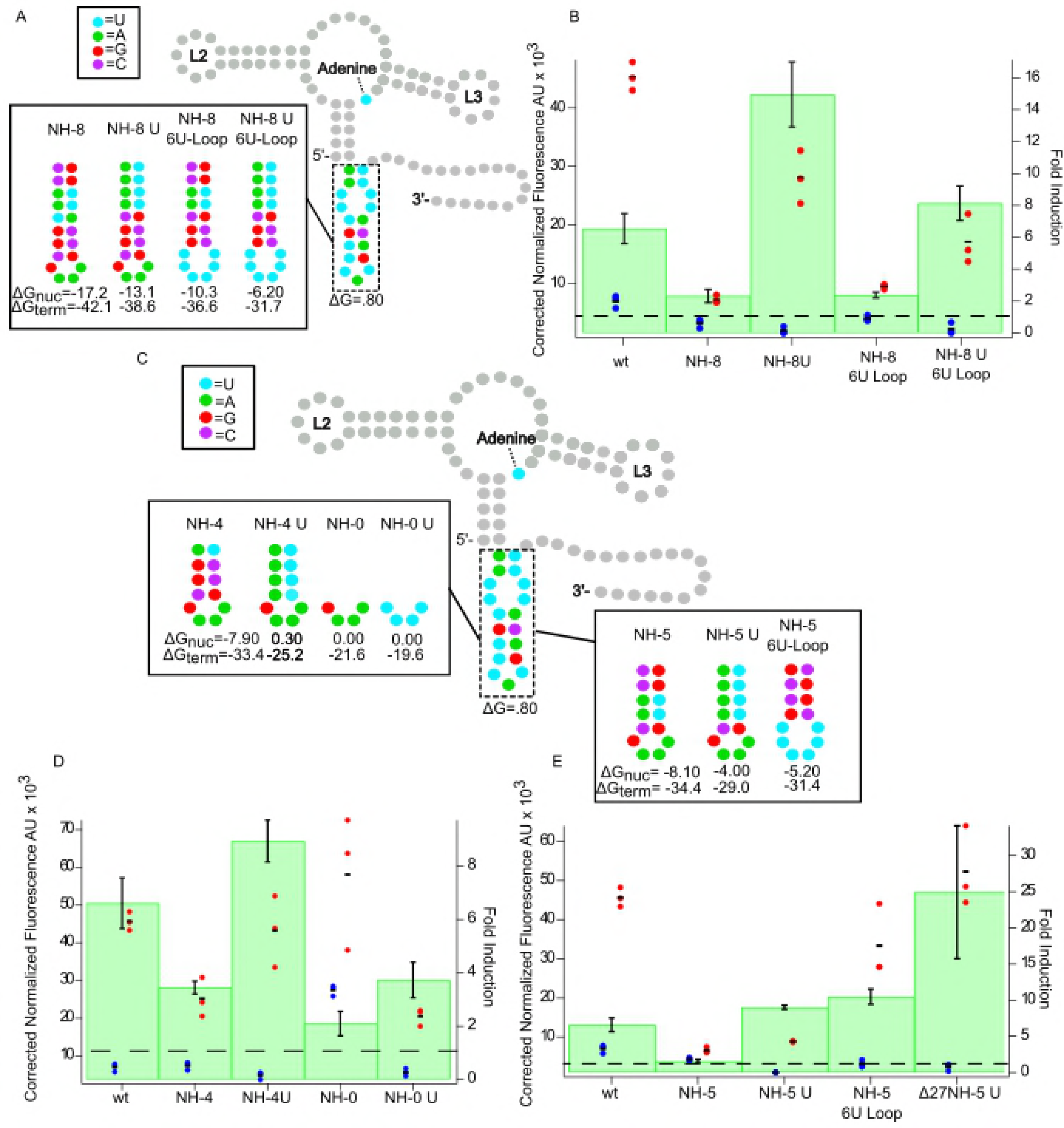
The hexa-uridine tract is necessary for efficient regulatory activity. (A) Secondary structure of the *pbuE* riboswitch highlighting each NH-8 mutant. The aptamer domain remains unchanged. The leader sequence has been omitted for simplicity. The nucleotide coloring scheme is the same as in Figure 1. (B) Background corrected normalized fluorescence data for each NH-8 mutant in the absence (blue) and presence (red) of 2-aminopurine. Each data point represents the average of three technical replicates of one biological replicate and the black bar represents fold induction of the average of the biological replicates. pBR327 is a negative control. Wild type data used for this figure collected on the same days as the other data in the figure. (C) Secondary structure of the *pbuE* riboswitch highlighting each NH-4, NH-0, and NH-5 mutants. (D) Background corrected normalized fluorescence data for each NH-4 and NH-0 mutant. (E) Background corrected normalized fluorescence data for each NH-5 mutant.

In an attempt to assimilate the various advantageous regulatory features observed in this study, a riboswitch was designed with no pre-aptamer sequence and a five base pair nucleator helix with a six-uridine pause on the 3’ side. This construct out-performed wild type and all other variants tested in this study with a fold induction near 25-fold (Fig 6E). The ability to pare down the expression platform to only the required elements with great success further reinforces the validity in approaches that exploit natural expression platforms are hosts for alternative aptamers to create novel riboswitches.

## Discussion

While the structure, conformational dynamics and ligand-binding properties of isolated riboswitch aptamer domains have been extensively studied, how these properties direct the expression platform to affect gene regulation remains poorly understood. In this current study, we have used a structure-guided mutagenesis strategy in concert with a cell-based fluorescent protein reporter to examine how sequence elements in both the aptamer and the expression platform influence the regulatory activity of the *B. subtilis pbuE* adenine riboswitch. These data reveal that many sequence elements outside of the core ligand-binding site and secondary structural switch play a significant role in establishing ligand-dependent reporter gene expression.

### Base pairs proximal to interhelical regions play a significant role in establishing regulatory activity

Analysis of critical functional regions of riboswitch aptamers have focused primarily on the first or second shell of nucleotides surrounding the ligand and those directly involved in establishing tertiary structure, which are almost invariably the most conserved nucleotides. In the purine riboswitch family, these nucleotides are found in the ligand-binding three-way junction and the terminal loop-loop interaction that establishes the helical packing of the aptamer. Between the *B. subtilis pbuE* and *xpt* aptamers, there are only 3 differences out of 27 nucleotides in J1/2, L2, J2/3, L3 and J3/1. Of these three, one is the specificity pyrimidine (*pbuE* nt 82) that determines adenine or guanine binding, one is a nucleotide that is flipped out into solution in the bound form of the riboswitch (*pbuE* nt 56) and one is a nucleotide in L2 that stacks but does not form any base-base interactions (*pbuE* nt 40). Thus, grafting the J1/2 through J3/1 region of *xpt*(C74U) onto *pbuE* with the compensatory base pair changes in the expression platform can be considered a very conservative aptamer exchange. The observation that the *xpt*(C74U)/*pbuE* aptamer/expresson platform chimera was non-functional in the cellular context indicates that there are unrealized features of the aptamer domain that are crucial for ligand-dependent secondary structural switching.

Unexpectedly, establishing regulatory function to the *xpt*(C74U)/*pbuE* chimera required two point-substitutions that are located ∼21 Å from each other in the structure of the *xpt* aptamer. Individually, each point substitution does not rescue activity, suggesting that these the point substitutions are required to either suppress two distinct defects in the aptamer domain or re-establish cross-talk between the loop-loop interaction and the three-way junction. If the latter is the case, this reveals that the conformational dynamics of the three-way junction and the L2-L3 interaction are interlinked. The linkage between L2-L3 dynamics and adenine binding has been previously observed in single-molecule and fluorescence lifetime measurements of the *Vibrio vulnificus add* adenine riboswitch [27-29]. As the importance of the L3 closing pair and the P2 tunebox on regulatory activity was further validated using chimeras with *B. subtilis yxjA*(C74U) and *purE*(C74U) aptamer domains, we propose that this linkage is a universal and essential feature of guanine and adenine riboswitches. Furthermore, it is very likely that since the wild type xpt(C74U) aptamer binds 2AP with high affinity as established in previous work [43] these point substitutions are required to restore some aspect of the linkage between ligand binding and the regulatory switch. This opens the intriguing possibility that aptamer dynamics in the apo state are not only crucial for ligand binding but also for efficient structural switching.

It is important to note that these restorative point substitutions reside not in non-canonical features but within adjacent helical elements. This reveals that the sequence elements at the termini of the helical elements play a significant role in RNA function. In an analysis of the base pairing patterns in the 50S ribosomal RNA, it was observed that the ends of helices are enriched in non-canonical pairs over the interior of helices (6.3% versus 1.1%) [52]. Base pairs in helices flanking junctions were found to play an important role in both local and global conformational dynamics of RNA [53]. In a recent example, the two base pairs flanking an internal bulge motif were found to influence ligand selectivity by a cobalamin riboswitch [54]. Notably, one of these pairs is a non-canonical A-C pair, as observed in the ligand-bound crystal structure [55]. Together, these data suggest that helical pairs—Watson-Crick or non-canonical—flanking interhelical regions should be considered as part of the junction or loop module. For example, the GNRA and UNYG tetraloop motifs are actually two nucleotide loops flanked by a non-canonical base pair (G-A or U-G, respectively) [56]. Further, the proximal Watson-Crick pair to tetraloop is often a C-G pair and is considered to be part of the extended tetraloop element and serves to impart significant thermostability to the loop [57]. This has implications for design of novel RNAs using a modular design approach with recurrent tertiary structural motifs [58]. Rather than considering just the module such as a terminal loop (for example, a tetraloop or T-loop) or internal loop (such as a kink-turn), flanking helical regions should be considered as part of the module.

### Efficient strand exchange is strongly dependent on the sequence of the P1 helix

Another helical structural element whose sequence composition has been proposed to be important for efficient ligand-dependent regulatory switching is the P1 helix. While the two base pairs proximal to the three-way junction are critical for ligand binding due to long-range interactions with bases in J2/3, the other base pairs in P1 are not directly involved in ligand recognition. Yet, phylogenetic alignments of purine riboswitches reveal a marked preference for either specific base pairs or R-Y versus Y-R orientation [19, 59]. Since these base pairs do not participate directly in ligand binding or ligand-induced RNA structure, the most likely explanation for sequence preferences in this region is to facilitate strand exchange between the P1 helix and an alternative helix in the expression platform.

The initial set of P1 sequence variants designed to test this hypothesis revealed no clear correlation between sequence and regulatory activity (Fig. 3B). A cotranscriptional analysis of several variants suggested that misfolding involving the pre-aptamer leader sequence and the aptamer can occur. This was validated for one variant (P1-GC2b) in which mutations introduced into the pre-aptamer sequence that disrupt this alternative fold rescue regulatory activity. Alternative structures involving the pre-aptamer sequence have been observed in the *V. vulnificus add* adenine riboswitch in which alternative secondary structure was proposed to be a critical feature of regulation at different temperatures [60]. To resolve this issue, the pre-aptamer leader sequence, which has been previously shown to have only slight effects on performance of the switch [33], was completely removed. These observations reinforce that the pre-aptamer sequence, while generally ignored, can nonetheless influence the regulatory activity of a riboswitch.

The sequence composition of the three base pairs at the 5’-side of P1 helix has a very strong influence over secondary structural switching. Surprisingly, conversion of the G-C pair to an A-U strongly impaired the ability of the terminator to form, even in the absence of 2AP, while three G-C pairs drove strong termination both in the presence and absence of 2AP. This suggests that the progress of strand exchange through the P1 helix by the competing terminator helix is less efficient in the context of weaker A-U pairs than with stronger G-C pairs. In contrast, robust regulatory activity is supported by either one or two G-C pairs in P1. Together, these results support a hypothesis that sequence conservation patterns in the P1 helix of many riboswitches are directly related to facilitating a rapid strand exchange process, a key component of the kinetically-controlled regulatory mechanism.

### Nucleation of the intrinsic terminator

Many riboswitches have secondary structural elements in the expression platform whose formation is independent of ligand binding to the aptamer. These elements presumably facilitate the switching function of the expression platform, such as by promoting rapid nucleation of alternative secondary structure. These features have previously been shown to be required for efficient regulatory activity in the lysine [61] and a purine riboswitch [33]. In the later study on the *pbuE* riboswitch, a limited set of variants that strengthened the terminator helix was tested, demonstrating that stabilizing the wild type nucleator element facilitated termination.

In this study, we designed a more systematic set of nucleation elements that explored both length and sequence composition of the nucleator. Unexpectedly, the length of the stem of the nucleator hairpin had only a moderate effect on the degree of activation of reporter expression. The stem length of the nucleator helix is clearly not the only structural feature that in the expression platform that optimizes performance. An element as short as four base pairs support near wild type levels of regulatory activity when a hexa-uridine tract is included in or directly following the antiterminator helix. However, even the absence of a helix supported minimal upregulation indicating that the presence of the nucleator is an important, but not essential, aspect of ligand-dependent secondary structural switching.

These data strongly support the role of a poly-uridine RNA polymerase pause site as a significant component of the regulatory switch. This finding is best exemplified by comparison of the NH-8 and NH-8 U variants. While an eight base pair nucleator helix shows strong switching performance (NH-8 U), conversion of two A-U pairs to G-C pairs that disrupt the 3’-uridine tract resulted in an almost complete loss of activity despite further thermodynamically stabilizing the intrinsic terminator. The same effect is observed with a five base pair nucleator helix as little to no switching activity occurs in the absence of a six uridine pause yet activity is fully recovered upon inclusion of the pause. Furthermore, all nucleator helix variants that preserve the stretch of uridine residues on the 3’ side of the helix showed strong ligand-dependent activation of reporter expression as seen in variants NH-8 U, NH-4 U, and NH-0 U.

The role of the pause sequence appears to be to increase the time window for ligand binding rather than nucleation of the terminator helix. In the wild type *pbuE* expression platform, the hexa-uridine sequence occurs at site where it could affect either the timeframe of ligand binding or nucleation of the terminator helix. Since hairpin nucleation is very rapid—on the order of microseconds [62, 63]—it is likely that the role of this pause is in increasing the binding time for the aptamer, as has been proposed for the *B. subtilis ribD* FMN riboswitch [13]. This is supported by the observation that the uridine tract could be moved to the loop of the nucleator helix where it can only serve to increase the time window for aptamer folding and ligand binding and still support regulatory activity. This indicates also that the precise positioning of poly-uridine tracts is not essential in the expression platform, as long as it is in a position that lengthens the timeframe of ligand binding prior to the onset of the strand invasion process.

### Prospects for the design of artificial riboswitches

The ability to engineer synthetic riboswitches that function in cells open doors for many applications. Of particular interest is the use of riboswitches as biosensors [64] that can sense the presence of certain molecules and provide a visual readout to indicate their presence. By knowing the requirements for optimal riboswitch performance, specificity for a particular ligand can be easily toggled while expression can be tuned precisely. Another application of interest rests in engineering genetic pathways in organisms that perform beneficial reactions. For example, chimeric riboswitches using parts derived from natural riboswitches have been used to establish an inducible genetic pathway in nitrogen fixing cyanobacteria, suggesting promise for controlling the varying abilities of cyanobacteria as well as other simple organisms [65].

While seemingly simple to design, artificial riboswitches are difficult to implement, particularly in the co-transcriptional and cellular context. In part, this is due to features of natural riboswitches beyond the aptamer and alternative secondary structure that contribute to efficient ligand-dependent regulation. In this work, we have revealed several features of the natural *pbuE* riboswitch that would be difficult to rationally consider in design using current tools. For example, predicting P1 helix sequences that promote strand exchange on timescales compatible with the rates of ligand binding to the aptamer domain cannot be computationally predicted. Further, rational design can be easily frustrated by alternative structures involving sequences outside of the core riboswitch making *de novo* design more difficult. Finally, conformational dynamics of the unbound state of the aptamer that influence rates of ligand binding and/or strand exchange cannot be accurately predicted or modeled. Adding to the above difficulties, the unanticipated interdependence of these elements that can hamper rational design.

Instead of a pure design approach, this work suggests that a hybrid design and screening approach might be more successful in implementing novel riboswitches. In this approach, an “approximate” riboswitch is created using computational design strategies and “mix-and-match” aptamer domains and expression platforms. In a second step, an interdomain sequence or communication module is randomized and a library of variants expressed in cells is screened for individual sequences with optimal properties, as in the “dual-selection” approach [66]. However, this study reveals that multiple regions of the riboswitch may have to be screened to fully optimize performance in cells—merely optimizing a communication element that couples the aptamer domain to a structural switch is likely not sufficient to generate a robust regulatory device.

## ACKNOWLEDGEMENTS

The authors would like to thank Drs. Jacob Polaski and Joan Marcano-Velazquez for helpful advice throughout this project.

## FUNDING

This work has been funded by a grant from the National Institutes of Health (R01 GM073850) to R.T.B.

## Supporting information

**S1 Table. Riboswitch mutant sequences**. ^a^ Full promoter and leader of wild type pbuE sequence through the polyuridine tract. The promoter is underlined and italicized, the leader sequence is in bold. ^b^ Full leader sequence in bold with mutations underlined of each of the remaining riboswitch mutants.

**S2 Table. Raw data values.** ^*^Data recollected in the context of the same cell prep as the constructs to which the data is being compared, used in Figure 4. ^**^Data recollected, used in Figure 6.

**S3 Figure. *pbuE* controls.** Direct comparison of pbuE full length, pbuE with first 11 nucleotides removed from the pre-aptamer sequence, and pbuE with 11 nucleotides removed and additional AatII and SpeI restriction sites added (RS). Each control was assayed in the absence (blue) and presence (red) of 2AP with the fold induction reported in green.

**S4 Figure. Structure of *purE*/*pbuE* hybrid with highlighted repair mutations.** Nucleotide coloring scheme is the same as Fig. 1. Pre-aptamer leader sequence removed for simplicity.

**S5 Figure. Structure of yxjA/pbuE hybrid with highlighted repair mutations.** Nucleotide coloring scheme is the same as for Fig. 1. Pre-aptamer leader sequence removed for simplicity.

## REFERENCES

1. Garst AD, Edwards AL, Batey RT. Riboswitches: structures and mechanisms. Cold Spring Harb Perspect Biol. 2011;3(6). Epub 2011/06/01. doi: 10.1101/cshperspect.a003533. PubMed PMID: 20943759; PubMed Central PMCID: PMCPMC3098680.

2. Ferré-D’Amaré AR, Scott WG. Small self-cleaving ribozymes. Cold Spring Harb Perspect Biol. 2010;2:a003574. Epub 2010/09/15. doi: 10.1101/cshperspect.a003574. PubMed PMID: 20843979; PubMed Central PMCID: PMCPMC2944367.

3. Breaker RR. Riboswitches and the RNA world. Cold Spring Harb Perspect Biol. 2012;4(2). Epub 2012/02/01. doi: 10.1101/cshperspect.a003566. PubMed PMID: 21106649; PubMed Central PMCID: PMCPMC3281570.

4. Bastet L, Dubé A, Massé E, Lafontaine DA. New insights into riboswitch regulation mechanisms. Mol Microbiol. 2011;80(5):1148–54. Epub 2011/04/20. doi: 10.1111/j.1365-2958.2011.07654.x. PubMed PMID: 21477128.

5. Wachter A. Gene regulation by structured mRNA elements. Trends Genet. 2014;30(5):172–81. Epub 2014/04/26. doi: 10.1016/j.tig.2014.03.001. PubMed PMID: 24780087..

6. Mellin JR, Cossart P. Unexpected versatility in bacterial riboswitches. Trends Genet. 2015;31(3):150–6. Epub 2015/02/21. doi: 10.1016/j.tig.2015.01.005. PubMed PMID: 25708284.

7. Berens C, Groher F, Suess B. RNA aptamers as genetic control devices: the potential of riboswitches as synthetic elements for regulating gene expression. Biotechnol J. 2015;10(2):246–57. doi: 10.1002/biot.201300498. PubMed PMID: 25676052.

8. Hallberg ZF, Su Y, Kitto RZ, Hammond MC. Engineering and In Vivo Applications of Riboswitches. Annu Rev Biochem. 2017;86:515–39. Epub 2017/03/30. doi: 10.1146/annurev-biochem-060815-014628. PubMed PMID: 28375743.

9. Findeiß S, Wachsmuth M, Mörl M, Stadler PF. Design of transcription regulating riboswitches. Methods Enzymol. 2015;550:1–22. Epub 2014/12/26. doi: 10.1016/bs.mie.2014.10.029. PubMed PMID: 25605378.

10. Etzel M, Mörl M. Synthetic Riboswitches: From Plug and Pray toward Plug and Play. Biochemistry. 2017;56(9):1181–98. Epub 2017/02/24. doi: 10.1021/acs.biochem.6b01218. PubMed PMID: 28206750.

11. Ceres P, Garst AD, Marcano-Velázquez JG, Batey RT. Modularity of select riboswitch expression platforms enables facile engineering of novel genetic regulatory devices. ACS Synth Biol. 2013;2(8):463–72. Epub 2013/03/28. doi: 10.1021/sb4000096. PubMed PMID: 23654267; PubMed Central PMCID: PMCPMC3742664.

12. Wickiser JK, Cheah MT, Breaker RR, Crothers DM. The kinetics of ligand binding by an adenine-sensing riboswitch. Biochemistry. 2005;44(40):13404–14. doi: 10.1021/bi051008u. PubMed PMID: 16201765.

13. Wickiser JK, Winkler WC, Breaker RR, Crothers DM. The speed of RNA transcription and metabolite binding kinetics operate an FMN riboswitch. Mol Cell. 2005;18(1):49–60. doi: 10.1016/j.molcel.2005.02.032. PubMed PMID: 15808508.

14. Garst AD, Porter EB, Batey RT. Insights into the regulatory landscape of the lysine riboswitch. J Mol Biol. 2012;423(1):17–33. Epub 2012/07/03. doi: 10.1016/j.jmb.2012.06.038. PubMed PMID: 22771573; PubMed Central PMCID: PMCPMC3444622.

15. Aboul-ela F, Huang W, Abd Elrahman M, Boyapati V, Li P. Linking aptamer-ligand binding and expression platform folding in riboswitches: prospects for mechanistic modeling and design. Wiley Interdiscip Rev RNA. 2015;6(6):631–50. Epub 2015/09/11. doi: 10.1002/wrna.1300. PubMed PMID: 26361734; PubMed Central PMCID: PMCPMC5049679.

16. Bastet L, Chauvier A, Singh N, Lussier A, Lamontagne AM, Prévost K, et al. Translational control and Rho-dependent transcription termination are intimately linked in riboswitch regulation. Nucleic Acids Res. 2017;45(12):7474–86. doi: 10.1093/nar/gkx434. PubMed PMID: 28520932; PubMed Central PMCID: PMCPMC5499598.

17. Chauvier A, Picard-Jean F, Berger-Dancause JC, Bastet L, Naghdi MR, Dubé A, et al. Transcriptional pausing at the translation start site operates as a critical checkpoint for riboswitch regulation. Nat Commun. 2017;8:13892. Epub 2017/01/10. doi: 10.1038/ncomms13892. PubMed PMID: 28071751; PubMed Central PMCID: PMCPMC5234074.

18. Proshkin S, Mironov A, Nudler E. Riboswitches in regulation of Rho-dependent transcription termination. Biochim Biophys Acta. 2014;1839(10):974–7. Epub 2014/04/13. doi: 10.1016/j.bbagrm.2014.04.002. PubMed PMID: 24731855.

19. McCown PJ, Corbino KA, Stav S, Sherlock ME, Breaker RR. Riboswitch diversity and distribution. RNA. 2017;23(7):995–1011. Epub 2017/04/10. doi: 10.1261/rna.061234.117. PubMed PMID: 28396576; PubMed Central PMCID: PMCPMC5473149.

20. Mandal M, Breaker RR. Adenine riboswitches and gene activation by disruption of a transcription terminator. Nat Struct Mol Biol. 2004;11(1):29–35. Epub 2003/12/29. doi: 10.1038/nsmb710. PubMed PMID: 14718920.

21. Porter EB, Marcano-Velázquez JG, Batey RT. The purine riboswitch as a model system for exploring RNA biology and chemistry. Biochim Biophys Acta. 2014;1839(10):919–30. Epub 2014/02/28. doi: 10.1016/j.bbagrm.2014.02.014. PubMed PMID: 24590258; PubMed Central PMCID: PMCPMC4148472.

22. Batey RT, Gilbert SD, Montange RK. Structure of a natural guanine-responsive riboswitch complexed with the metabolite hypoxanthine. Nature. 2004;432(7015):411–5. doi: 10.1038/nature03037. PubMed PMID: 15549109.

23. Serganov A, Yuan YR, Pikovskaya O, Polonskaia A, Malinina L, Phan AT, et al. Structural basis for discriminative regulation of gene expression by adenine- and guanine-sensing mRNAs. Chem Biol. 2004;11(12):1729–41. doi: 10.1016/j.chembiol.2004.11.018. PubMed PMID: 15610857; PubMed Central PMCID: PMCPMC4692365.

24. Chandra V, Hannan Z, Xu H, Mandal M. Single-molecule analysis reveals multi-state folding of a guanine riboswitch. Nat Chem Biol. 2017;13(2):194–201. Epub 2016/12/12. doi: 10.1038/nchembio.2252. PubMed PMID: 27941758.

25. Brenner MD, Scanlan MS, Nahas MK, Ha T, Silverman SK. Multivector fluorescence analysis of the xpt guanine riboswitch aptamer domain and the conformational role of guanine. Biochemistry. 2010;49(8):1596–605. doi: 10.1021/bi9019912. PubMed PMID: 20108980; PubMed Central PMCID: PMCPMC2854158.

26. Noeske J, Buck J, Fürtig B, Nasiri HR, Schwalbe H, Wöhnert J. Interplay of ‘induced fit’ and preorganization in the ligand induced folding of the aptamer domain of the guanine binding riboswitch. Nucleic Acids Res. 2007;35(2):572–83. Epub 2006/12/14. doi: 10.1093/nar/gkl1094. PubMed PMID: 17175531; PubMed Central PMCID: PMCPMC1802621.

27. Lemay JF, Penedo JC, Tremblay R, Lilley DM, Lafontaine DA. Folding of the adenine riboswitch. Chem Biol. 2006;13(8):857–68. doi: 10.1016/j.chembiol.2006.06.010. PubMed PMID: 16931335.

28. Eskandari S, Prychyna O, Leung J, Avdic D, O’Neill MA. Ligand-directed dynamics of adenine riboswitch conformers. J Am Chem Soc. 2007;129(37):11308–9. Epub 2007/08/22. doi: 10.1021/ja073159l. PubMed PMID: 17713907.

29. Prychyna O, Dahabieh MS, Chao J, O’Neill MA. Sequence-dependent folding and unfolding of ligand-bound purine riboswitches. Biopolymers. 2009;91(11):953–65. doi: 10.1002/bip.21283. PubMed PMID: 19603494.

30. Lemay JF, Desnoyers G, Blouin S, Heppell B, Bastet L, St-Pierre P, et al. Comparative Study between Transcriptionally- and Translationally-Acting Adenine Riboswitches Reveals Key Differences in Riboswitch Regulatory Mechanisms. PLoS Genet. 2011;7:1. doi: 10.1371/journal.pgen.1001278. PubMed PMID: WOS:000286653500017.

31. Frieda KL, Block SM. Direct observation of cotranscriptional folding in an adenine riboswitch. Science. 2012;338(6105):397–400. doi: 10.1126/science.1225722. PubMed PMID: 23087247; PubMed Central PMCID: PMCPMC3496414.

32. Rieder R, Lang K, Graber D, Micura R. Ligand-induced folding of the adenosine deaminase A-riboswitch and implications on riboswitch translational control. Chembiochem. 2007;8(8):896–902. doi: 10.1002/cbic.200700057. PubMed PMID: 17440909.

33. Marcano-Velázquez JG, Batey RT. Structure-guided mutational analysis of gene regulation by the Bacillus subtilis pbuE adenine-responsive riboswitch in a cellular context. J Biol Chem. 2015;290(7):4464–75. Epub 2014/12/30. doi: 10.1074/jbc.M114.613497. PubMed PMID: 25550163; PubMed Central PMCID: PMCPMC4326850.

34. Prodromou C, Pearl LH. Recursive PCR: a novel technique for total gene synthesis. Protein Eng. 1992;5(8):827–9. PubMed PMID: 1287665.

35. Sambrook J, Russell DW. Molecular cloning : a laboratory manual. 3rd ed. Cold Spring Harbor, N.Y.: Cold Spring Harbor Laboratory Press; 2001.

36. Ceres P, Trausch JJ, Batey RT. Engineering modular ‘ON’ RNA switches using biological components. Nucleic Acids Res. 2013;41(22):10449–61. Epub 2013/09/02. doi: 10.1093/nar/gkt787. PubMed PMID: 23999097; PubMed Central PMCID: PMCPMC3905868.

37. Davis JH, Rubin AJ, Sauer RT. Design, construction and characterization of a set of insulated bacterial promoters. Nucleic Acids Res. 2011;39(3):1131–41. Epub 2010/09/17. doi: 10.1093/nar/gkq810. PubMed PMID: 20843779; PubMed Central PMCID: PMCPMC3035448.

38. Baba T, Ara T, Hasegawa M, Takai Y, Okumura Y, Baba M, et al. Construction of Escherichia coli K-12 in-frame, single-gene knockout mutants: the Keio collection. Mol Syst Biol. 2006;2:2006.0008. Epub 2006/02/21. doi: 10.1038/msb4100050. PubMed PMID: 16738554; PubMed Central PMCID: PMCPMC1681482.

39. Xayaphoummine A, Bucher T, Isambert H. Kinefold web server for RNA/DNA folding path and structure prediction including pseudoknots and knots. Nucleic Acids Res. 2005;33(Web Server issue):W605–10. doi: 10.1093/nar/gki447. PubMed PMID: 15980546; PubMed Central PMCID: PMCPMC1160208.

40. Stoddard CD, Batey RT. Mix-and-match riboswitches. ACS Chem Biol. 2006;1(12):751–4. doi: 10.1021/cb600458w. PubMed PMID: 17240972.

41. Gilbert SD, Stoddard CD, Wise SJ, Batey RT. Thermodynamic and kinetic characterization of ligand binding to the purine riboswitch aptamer domain. J Mol Biol. 2006;359(3):754–68. Epub 2006/04/21. doi: 10.1016/j.jmb.2006.04.003. PubMed PMID: 16650860.

42. Gilbert SD, Love CE, Edwards AL, Batey RT. Mutational analysis of the purine riboswitch aptamer domain. Biochemistry. 2007;46(46):13297–309. Epub 2007/10/26. doi: 10.1021/bi700410g. PubMed PMID: 17960911; PubMed Central PMCID: PMCPMC2556308.

43. Stoddard CD, Widmann J, Trausch JJ, Marcano-Velázquez JG, Knight R, Batey RT. Nucleotides adjacent to the ligand-binding pocket are linked to activity tuning in the purine riboswitch. J Mol Biol. 2013;425(10):1596–611. Epub 2013/02/26. doi: 10.1016/j.jmb.2013.02.023. PubMed PMID: 23485418; PubMed Central PMCID: PMCPMC3769107.

44. Mulhbacher J, Lafontaine DA. Ligand recognition determinants of guanine riboswitches. Nucleic Acids Res. 2007;35(16):5568–80. Epub 2007/08/17. doi: 10.1093/nar/gkm572. PubMed PMID: 17704135; PubMed Central PMCID: PMCPMC2018637.

45. Greenleaf WJ, Frieda KL, Foster DA, Woodside MT, Block SM. Direct observation of hierarchical folding in single riboswitch aptamers. Science. 2008;319(5863):630–3. Epub 2008/01/03. doi: 10.1126/science.1151298. PubMed PMID: 18174398; PubMed Central PMCID: PMCPMC2640945.

46. Blouin S, Chinnappan R, Lafontaine DA. Folding of the lysine riboswitch: importance of peripheral elements for transcriptional regulation. Nucleic Acids Res. 2011;39(8):3373–87. Epub 2010/12/17. doi: 10.1093/nar/gkq1247. PubMed PMID: 21169337; PubMed Central PMCID: PMCPMC3082890.

47. Zhang DY, Winfree E. Control of DNA strand displacement kinetics using toehold exchange. J Am Chem Soc. 2009;131(47):17303–14. doi: 10.1021/ja906987s. PubMed PMID: 19894722.

48. Srinivas N, Ouldridge TE, Sulc P, Schaeffer JM, Yurke B, Louis AA, et al. On the biophysics and kinetics of toehold-mediated DNA strand displacement. Nucleic Acids Res. 2013;41(22):10641–58. Epub 2013/09/09. doi: 10.1093/nar/gkt801. PubMed PMID: 24019238; PubMed Central PMCID: PMCPMC3905871.

49. Perdrizet GA, Artsimovitch I, Furman R, Sosnick TR, Pan T. Transcriptional pausing coordinates folding of the aptamer domain and the expression platform of a riboswitch. Proc Natl Acad Sci U S A. 2012;109(9):3323–8. Epub 2012/02/13. doi: 10.1073/pnas.1113086109. PubMed PMID: 22331895; PubMed Central PMCID: PMCPMC3295289.

50. Steinert H, Sochor F, Wacker A, Buck J, Helmling C, Hiller F, et al. Pausing guides RNA folding to populate transiently stable RNA structures for riboswitch-based transcription regulation. Elife. 2017;6. Epub 2017/05/25. doi: 10.7554/eLife.21297. PubMed PMID: 28541183; PubMed Central PMCID: PMCPMC5459577.

51. Larson MH, Mooney RA, Peters JM, Windgassen T, Nayak D, Gross CA, et al. A pause sequence enriched at translation start sites drives transcription dynamics in vivo. Science. 2014;344(6187):1042–7. Epub 2014/05/01. doi: 10.1126/science.1251871. PubMed PMID: 24789973; PubMed Central PMCID: PMCPMC4108260.

52. Lee JC, Gutell RR. Diversity of base-pair conformations and their occurrence in rRNA structure and RNA structural motifs. J Mol Biol. 2004;344(5):1225–49. doi: 10.1016/j.jmb.2004.09.072. PubMed PMID: 15561141.

53. Stelzer AC, Kratz JD, Zhang Q, Al-Hashimi HM. RNA dynamics by design: biasing ensembles towards the ligand-bound state. Angew Chem Int Ed Engl. 2010;49(33):5731–3. doi: 10.1002/anie.201000814. PubMed PMID: 20583015; PubMed Central PMCID: PMCPMC3319304.

54. Polaski JT, Webster SM, Johnson JE, Batey RT. Cobalamin riboswitches exhibit a broad range of ability to discriminate between methylcobalamin and adenosylcobalamin. J Biol Chem. 2017;292(28):11650–8. Epub 2017/05/08. doi: 10.1074/jbc.M117.787176. PubMed PMID: 28483920; PubMed Central PMCID: PMCPMC5512062.

55. Johnson JE, Reyes FE, Polaski JT, Batey RT. B12 cofactors directly stabilize an mRNA regulatory switch. Nature. 2012;492(7427):133–7. Epub 2012/10/14. doi: 10.1038/nature11607. PubMed PMID: 23064232; PubMed Central PMCID: PMCPMC3518761.

56. Thapar R, Denmon AP, Nikonowicz EP. Recognition modes of RNA tetraloops and tetraloop-like motifs by RNA-binding proteins. Wiley Interdiscip Rev RNA. 2014;5(1):49–67. Epub 2013/10/03. doi: 10.1002/wrna.1196. PubMed PMID: 24124096; PubMed Central PMCID: PMCPMC3867596.

57. Blose JM, Proctor DJ, Veeraraghavan N, Misra VK, Bevilacqua PC. Contribution of the closing base pair to exceptional stability in RNA tetraloops: roles for molecular mimicry and electrostatic factors. J Am Chem Soc. 2009;131(24):8474–84. doi: 10.1021/ja900065e. PubMed PMID: 19476351.

58. Grabow WW, Jaeger L. RNA Self-Assembly and RNA Nanotechnology. Accounts of Chemical Research. 2014;47(6):1871–80. doi: 10.1021/ar500076k.

59. Barrick JE, Breaker RR. The distributions, mechanisms, and structures of metabolite-binding riboswitches. Genome Biol. 2007;8:R239. doi: 10.1186/gb-2007-8-11-r239. PubMed PMID: 17997835; PubMed Central PMCID: PMCPMC2258182.

60. Reining A, Nozinovic S, Schlepckow K, Buhr F, Fürtig B, Schwalbe H. Three-state mechanism couples ligand and temperature sensing in riboswitches. Nature. 2013;499(7458):355–9. Epub 2013/07/10. doi: 10.1038/nature12378. PubMed PMID: 23842498.

61. Blouin S, Lafontaine DA. A loop loop interaction and a K-turn motif located in the lysine aptamer domain are important for the riboswitch gene regulation control. RNA. 2007;13(8):1256–67. Epub 2007/06/21. doi: 10.1261/rna.560307. PubMed PMID: 17585050; PubMed Central PMCID: PMCPMC1924893.

62. Zhang W, Chen SJ. RNA hairpin-folding kinetics. Proc Natl Acad Sci U S A. 2002;99(4):1931–6. Epub 2002/02/12. doi: 10.1073/pnas.032443099. PubMed PMID: 11842187; PubMed Central PMCID: PMCPMC122297.

63. Zhang W, Chen SJ. Exploring the complex folding kinetics of RNA hairpins: I. General folding kinetics analysis. Biophys J. 2006;90(3):765–77. Epub 2005/11/04. doi: 10.1529/biophysj.105.062935. PubMed PMID: 16272440; PubMed Central PMCID: PMCPMC1367102.

64. Findeiß S, Etzel M, Will S, Mörl M, Stadler PF. Design of Artificial Riboswitches as Biosensors. Sensors (Basel). 2017;17(9). Epub 2017/08/30. doi: 10.3390/s17091990. PubMed PMID: 28867802; PubMed Central PMCID: PMCPMC5621056.

65. Higo A, Isu A, Fukaya Y, Hisabori T. Designing Synthetic Flexible Gene Regulation Networks Using RNA Devices in Cyanobacteria. ACS Synth Biol. 2017;6(1):55–61. doi: 10.1021/acssynbio.6b00201. PubMed PMID: WOS:000392575700007.

66. Sinha J, Topp S, Gallivan JP. From SELEX to cell dual selections for synthetic riboswitches. Methods Enzymol. 2011;497:207–20. Epub 2011/05/24. doi: 10.1016/B978-0-12-385075-1.00009-3. PubMed PMID: 21601088.

